# Longitudinal conditional probability of symmetric DNA methylation in *Arabidopsis* plants

**DOI:** 10.1101/2023.12.28.573516

**Authors:** Humberto Fernandes

## Abstract

DNA replication in high eukaryotes requires the faithful transmission of both the sequence and its methylation status. DNA methyltransferases are not consistently reliable enzymes for accurately restoring methylation in newly synthesised DNA strands when working in isolation. In this study, we evaluated some of the factors that enhance the fidelity of DNA methyltransferases and measure their local and distant impacts on the neighbouring methylation status. A clear understanding of DNA methylation fidelity during replication is important for advances in epigenetic knowledge in general and the potential development of new diagnostics and therapies for epigenetic diseases. We used the *Arabidopsis* model system because it is moderately methylated, which is an important advantage over animal models since it reduces the chance of a fully-methylated symmetric methylation to virtually nil while providing sufficient occupancy to ensure reliable interpretations. In this study, we employed hairpin bisulfite and nanopore sequencing (1D and 1D^2^) to test the inter- and intra-strand relationships among the neighbouring 5-methylcytosines (5mC) and evaluate the distribution of hemimethylated sites, as well as the correlations to other cues such as DNA methylation and histone marks. We observed that the presence of a CpG methylation site can predict the occurrence of neighbouring CpG methylation sites, but not necessarily CWG sites, whereas CWG methylation can predict both neighbouring CWG and CpG methylation locations. These predictabilities extend far beyond the size of a nucleosome, and sites 4 kb away still show higher probabilities of methylation compared to the global average.

**Graphic abstract:** Graphic abstract
Fidelity and longitudinal conditional methylation probability of CpG and CWG DNA methylation in *Arabidopsis* plants

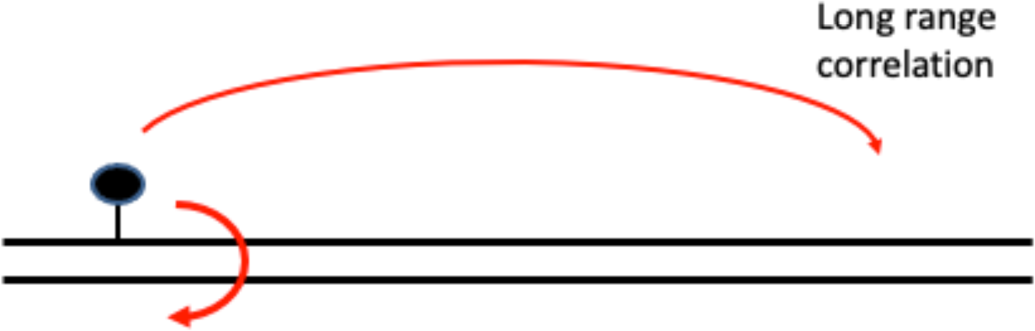

## Introduction

Faithful transmission of genetic information from parent to daughter cells requires accurate replication of the sequence content as well as reliable duplication of the DNA methylation state (Riggs & Xiong, 2004). The level of DNA methylation is determined through the combined actions of DNA methylation, which is capable of both maintenance and *de novo* activities, and demethylation enzymes. Maintenance, in particular, relies on the recruitment of DNA methyltransferases to the replication fork since DNA polymerases insert non-methylated cytosines during replication which may need to be methylated. Recently, it has been shown that replication-coupled DNA methylation does not fully account for this maintenance, as although it is achieved in 80% within 30 minutes of the fork passage, the remaining maintenance occurs over 10 hours through replication-uncoupled mechanisms (Ming et al., 2020). Excluding some well-documented dynamic changes during development, in both plants and mammals (B. P. Williams & Gehring, 2020), DNA methylation fidelity is a crucial process which, if established at the incorrect time and/or place or in an aberrant manner, can be linked to tumours and other diseases (Portela & Esteller, 2010; Ushijima et al., 2005). Early observations indicate that DNA methylases will scramble the original information in the first few rounds of replication without fidelity enhancement factors; therefore, other mechanisms involving feedback loops (B. P. Williams & Gehring, 2020), are utilised to increase DNA methylation fidelity and faithfully replicate the methylation patterns.

In high eukaryotes, the majority of DNA base modifications occur in the form of a 5-methylcytosine (5mC). The simplest and most commonly observed context involves symmetrical CpG dyads (^m^CG/G^m^C), whereby the requirement of faithful DNA methylation duplication does not occur because: DNA methyltransferases fail to methylate a cytosine on the daughter strand opposite to an ^m^CG on a newly synthesised strand, methylate a cytosine *de novo* without having a cue from the counter-strand (that was unmethylated), or when there is complete or partial (just on one strand) demethylase activity (K. Williams et al., 2011). In other words, methylation pattern maintenance must occur at both methylated and unmethylated sites (Ushijima et al., 2003). Mammalians exceptions to CpG methylation (non-CpG methylations) are observed at early stages of development and in brain tissue. A considerable number (∼1/10) of CpGs that remain hemimethylated in embryonic stem cells (ESCs) (Zhao et al., 2014) and trophoblastic stem cells (Sharif et al., 2016), and some of those hemimethylated CpG sites have recently been determined to be specific, persistent, inherited over several cell divisions and involved in regulating CTCF-mediated chromatin interactions (Xu & Corces, 2018). In addition to CpG, plants adopt CHG and CHH methylation activities, which involve typical maintenance as well as *de novo* methylation guided by histone modifications and RNA.

Overexpression of maintenance DNA methyltransferases results in methylation of previously unmethylated sites in both mammals (Biniszkiewicz et al., 2002; Takagi, Tajima, & Asano, 1995; Vertino, Yen, Gao, & Baylin, 1996), and plants (Brocklehurst et al., 2018). Recently, a transcriptional repressor (TCX5) was found in plants which regulates the expression of MET1 and CMT3 enzymes (Ning et al., 2020). TCX5 is part of the DREAM complex, which is conserved from plants to animals, and thus silencing of maintenance DNA methyltransferase genes may be a conserved feature in eukaryotes (Ning et al., 2020).

The deposition of 5mC epigenetic mark represses transposable elements and is traditionally associated with gene silencing; however, its more dynamic regulation role has been uncovered. DNA methylation is now considered important for embryonic development through the provision of cellular memory and modulation of gene regulation specificity in different cell types (Suelves, Carrio, Nunez-Alvarez, & Peinado, 2016). DNA methylation is concentrated in CpG islands, which modulate transcriptional activity by changing the binding capacities of transcriptional factors, positioning nucleosomes and linking methyl CpG binding proteins (Suelves et al., 2016). In addition, DNA methylation occurs in gene bodies where it promotes gene expression, although only (indirectly) by repressing their mechanics. DNA methylation prevents erogenous transcription in these structures, thereby releasing the machinery to the ‘true’ start site with the net effect of an increased expression of marked genes. Considering the need for stable transmission of methylation information, particularly during development and cell differentiation, a high fidelity of the DNA methylation system is required. In reality, clonal cells display epigenetic heterogeneity (with consequences on transcription variability, phenotype, drug response, and lineage specification) (Wang et al., 2020); however, this epigenetic drift is corrected over time and epigenome integrity is largely maintained. Flowering plants have a stable inheritance of DNA methylation across generations and show enhanced fidelity in sexual lineages compared to that of somatic cells, despite seemingly similar mechanisms (Hsieh et al., 2016; Park et al., 2016).

DNMT1 is a prototype enzyme for DNA maintenance methyltransferases (Law & Jacobsen, 2010). The enzyme has a preference for hemimethylated DNA (Gruenbaum et al., 1982; Nishiyama et al., 2016), although it cannot independently ensure the fidelity observed after replication and associates with PCNA and chromatin-associated protein UHRF1 in cells (Law & Jacobsen, 2010; Nishiyama et al., 2016; Rose & Klose, 2014). MET1 is the default DNA methyltransferase maintenance enzyme in plants (Law & Jacobsen, 2010) and it is, similar to its mammalian counterpart, connected to the replication fork through interactions with VIM proteins (Woo et al., 2008). Plants contain a larger number of maintenance DNA methyltransferases than mammals. They are in addition to MET1 equipped with CMT3, responsible for maintenance of CWG methylation, and CTM2 maintaining methylation of certain CHH sites. The remaining CHH methylation sites are preserved after replication by *de novo* methylation activity of the DRM2 enzyme (Law & Jacobsen, 2010; Stroud et al., 2014; Zemach et al., 2013). For the latter, a new CHH methylation single-read methylome data analysis allows to estimate the probability of each enzyme depositing a methyl group in a particular cysteine at an individual read, and to interrogate the variations among cells in heterogenous samples (Harris & Zemach, 2020).

Whole genome BS sequencing results that clearly ascertain variations in methylation patterns between molecules (Riggs & Xiong, 2004), as well as early and recent attempts to calculate DNA methylation fidelity (Harris, Lloyd, Domb, Zilberman, & Zemach, 2019; Minoguchi & Iba, 2008; Ushijima et al., 2003; Ushijima et al., 2005) have been accumulating; however, standard protocols currently only allow for the inference of this fidelity (Ushijima et al., 2003; Zhao et al., 2014). A limited number of genomic loci have been studied in detail with regard to the optics of fidelity (*i.e.* through the measurement of the two complementary strand methylation patterns of individual DNA molecules) (Arand et al., 2012; Burden et al., 2005; Laird et al., 2004; Xie et al., 2011), and genome-scale views are gradually emerging that couple BS with hairpin sequencing to generate accurate 5mC positioning in both strands (Ming et al., 2020; Wang et al., 2020; Xie et al., 2011). The paradox of expected clonal DNA methylation drift compared to observed stable propagation was recently addressed by Wang et al., who uncovered imprecise DNA maintenance activity of DNMT1, coupled with *de novo* activity of the same enzyme (Fatemi, Hermann, Gowher, & Jeltsch, 2002; Jeltsch & Jurkowska, 2014; Lorincz, Schubeler, Hutchinson, Dickerson, & Groudine, 2002), guided by neighbouring DNA methylation (Wang et al., 2020).

Some mechanisms of action of DNA methylation maintenance enzymes are well understood, but their fidelity has been understudied so far. Also, growing evidence indicates that the traditional site-specific DNA maintenance methylation model does not fully explain the theoretical models or experimental observations (Haerter, Lovkvist, Dodd, & Sneppen, 2014; Jeltsch & Jurkowska, 2014; Jones & Liang, 2009; Lovkvist, Dodd, Sneppen, & Haerter, 2016; Ming et al., 2020; Utsey & Keener, 2020; Wang et al., 2020). In this study, we used next-generation sequencing to determine which factors are involved in enhancing the fidelity of MET1, which is the DNA methyltransferase unit of plants. We used *Arabidopsis* because its DNA methyltransferases are involved in maintenance of contexts other than CpG and are viable, characterised, and available as unpaired mutants. In addition, global levels are ‘only’ moderately methylated in plants. Therefore, the accuracy for proper 5mC positioning is imperative and by virtue of its ‘scarcity’, the probability of obtaining a full-methylation CpG by chance is virtually nil (for mammals with generally high levels of methylation this assumption cannot be made), while a sufficient occupancy is provided to ensure the interpretations are reliable. Using hairpin BS-seq, we analysed the fidelity of MET1 and CMT3 and uncovered the probability of a longitudinal conditional methylation probability in *Arabidopsis* plants. The extension of this cannot be determined through single nucleosome wrapping as it occurs over 4 kb, as determined by nanopore long-read sequencing. In addition, we observed a strong correlation between CpG and CWG methylated sites and the methylation predictability of neighbouring CpG and CWG sites. Furthermore, CWG sites were shown to be strong predictors of neighbouring CpG sites, whereas the reverse was not observed.

## Results

To determine the level and distribution of hemimethylation in bulk *Arabidopsis* genomes, we performed whole-genome hairpin bisulfite sequencing (hairpin BS-seq) with 22-day-old WT *Arabidopsis* seedlings.

For the analysis, we used a robust algorithm which retained sequences only when the four sequences aligned (left and right from the hairpin and original and PCR-generated strands). As the protocol was BS-seq derived, we allowed C/T divergences that corresponded to the hemimethylated positions. However, each base still required the two reads (original and PCR amplified) to align (see methods). We then obtained robust hemimethylated sites for the WT *Arabidopsis* genome and divided the reads into two groups: sequences that mapped uniquely to the *Arabidopsis* genome (as a proxy for open chromatin) and those that mapped to two or more sites of the *Arabidopsis* genome (as a proxy for closed chromatin). For the first group (open chromatin), 10 and 1.5% of Cs were full methylations for CpG and CWG contexts, respectively, and 2.5 and 2.4% were hemimethylated for CpG and CWG, respectively. Regarding closed chromatin, the methylation numbers were higher, which was expected since 5mC is a mark of repressed chromatin, 32%/10% and 9%/7% for CpG/CWG full and hemimethylated sites, respectively. This approach of grouping the next generation reads into two categories (open and closed chromatin) was subsequently supported by nanopore reads copy number mapping to different chromosomes and organelles in which there was a clear increase of reads to the centromeres, assumed to be closed chromatin (see below).

### Conditional methylation probabilities in *Arabidopsis* WT plants

To investigate the probability of cytosine methylation in sites neighbouring a 5mC position, we plotted our bisulfite-converted sequence data and anchored the reads for each fully methylated CpG and CWG position. The meta-analysis shows that the CpG methylation typified that of a neighbour CpG that was fully methylated (Fig. 1A1), both with open and closed chromatins (Fig. 1A2). In addition, CWG was representative of a fully methylated neighbour CWG, although to a lesser extent (Fig. 1B). The data also indicated that CpG methylation was a poor predictor of a neighbouring methylated CWG (Fig. 1C), whereas a methylated CWG was a strong predictor of a neighbouring CpG methylation (Fig. 1D). The same exercise with anchoring in non-methylated Cs revealed that the effect was not random, as only unmethylated Cs were predicted (Fig. S1A-D).

**Figure 1:**
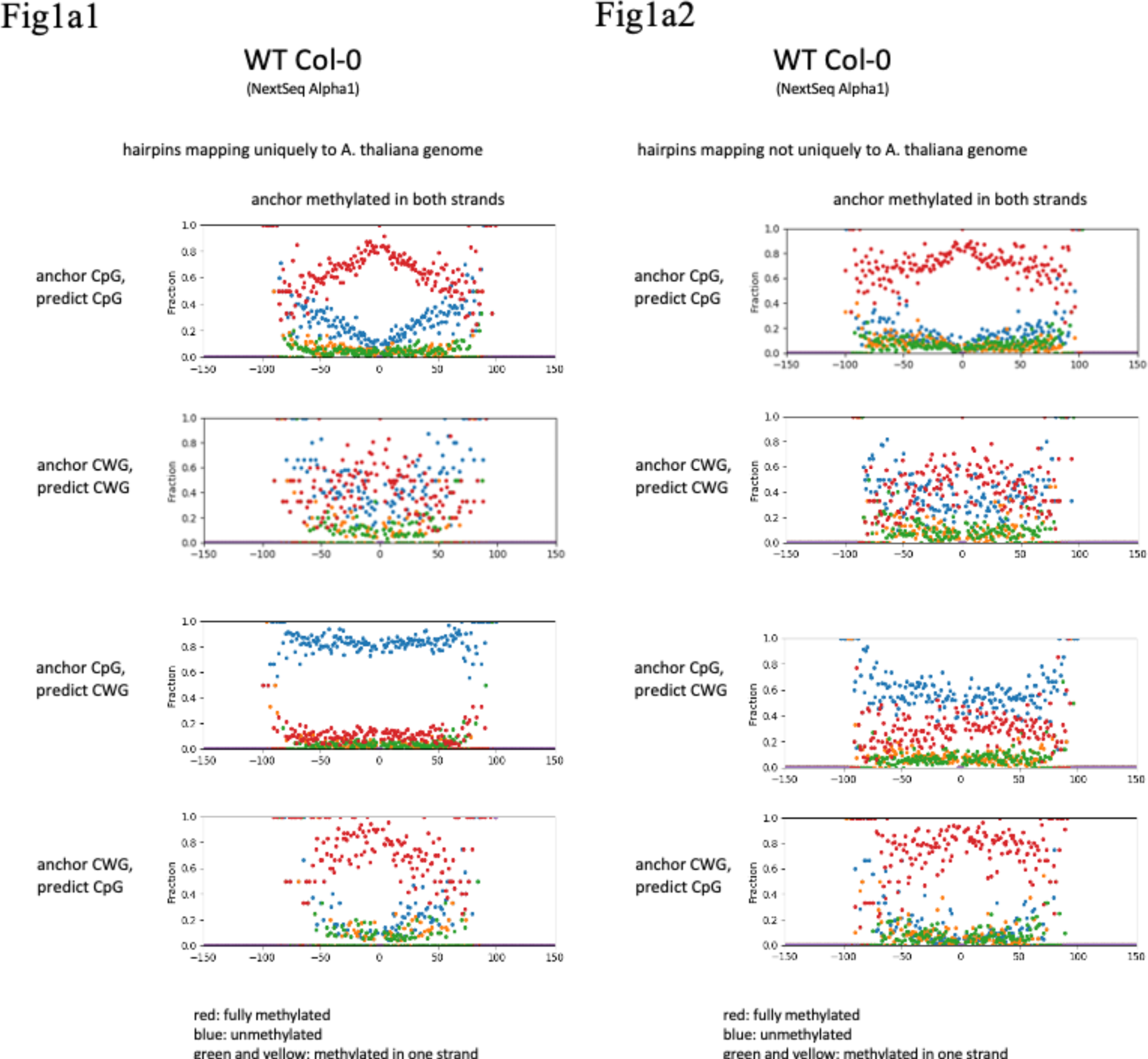
DNA methylation predictability of CpG and CWG neighbouring a fully-methylated CpG or CWG site in WT plants for reads that mapped uniquely (1a1) and not uniquely (1a2) to *Arabidopsis* genomic DNA. Red dots represent fully methylated sites, blue indicate unmethylated sites and green and yellow signify hemimethylated sites.

In addition, the metadata showed that although predictability of methylation faded with distance, it was still present at a single nucleosome interval, since the fraction of methylated Cs at the end of the plottable data was higher than the global average reported.

### Conditional methylation probabilities for several *Arabidopsis* mutants using NextSeq

To determine the influence of other epigenetic factors in the levels of conditional methylation probabilities, we performed the analysis described above using 11 *Arabidopsis* mutants with disturbed CpG methylation and overlapping characteristics that affect CpG, CHG and CHH methylation in this plant. In addition, we tested direct DNA disruption and indirect (via histone demethylation) DNA methylation disruption. To understand both DNA and trans-histone effects, we used *met1-3*, *hub1-5*, *atprmt10-1*, *vim1-2*, *vim2-2*, *drm1/drm2/kyp6*, *drm1/drm2/cmt3*, *drm1/drm2*, *kyp6*, *met1-7*, and *mat4* mutants. In combination, these mutants covered disruption of DNA methylation, histone methylation and ubiquitination at both read and write levels.

A decrease in CpG methylation levels of a heterozygote *met1-3* mutant (as homozygote *met1* mutant has too little CpG methylation to be relevant) was expected, which was observed in our analysis (Fig. 2), although this was not noted in relation to other mutants, as also confirmed (Fig. 2). However, the probability of a disturbance in neighbouring methylation in the *met1-3* mutant cannot be predicted without analysis. Meta-analysis of the heterozygote *met1-3* mutant produced a sharper decline in methylation probability around an anchor CpG compared to that of the WT (Fig. 3A). This level of decrease in methylation probability was not observed for mutants with directly impacted non-CpG methylation (such as mutants with impeded CWG and CHH methylation) (Figure 3B-L). CHG methylation was decreased when CMT3 or Kyp6 were disturbed (Fig. 2), either alone (*kyp6*) or in combination with *drm1*/*drm2* (*cmt3* and *kyp6*), which was expected since the two enzymes control CHG methylation in a feedback loop. Decrease of CWG methylation for such mutants is sufficient to abolish the predictability of a CpG methylation by neighbouring CWG methylation on not uniquely mapping reads (Fig. 3; right panels). The reduction of CHH methylation for mutants when DRM1 or DRM2 were disrupted was negligible because (i) our conversion rates were 0.3-1%, and (ii) CMT2 also plays a role in CHH methylation and is fully functional in these plants (perhaps even stimulated to compensate the losses of the DRM proteins).

**Figure 2:**
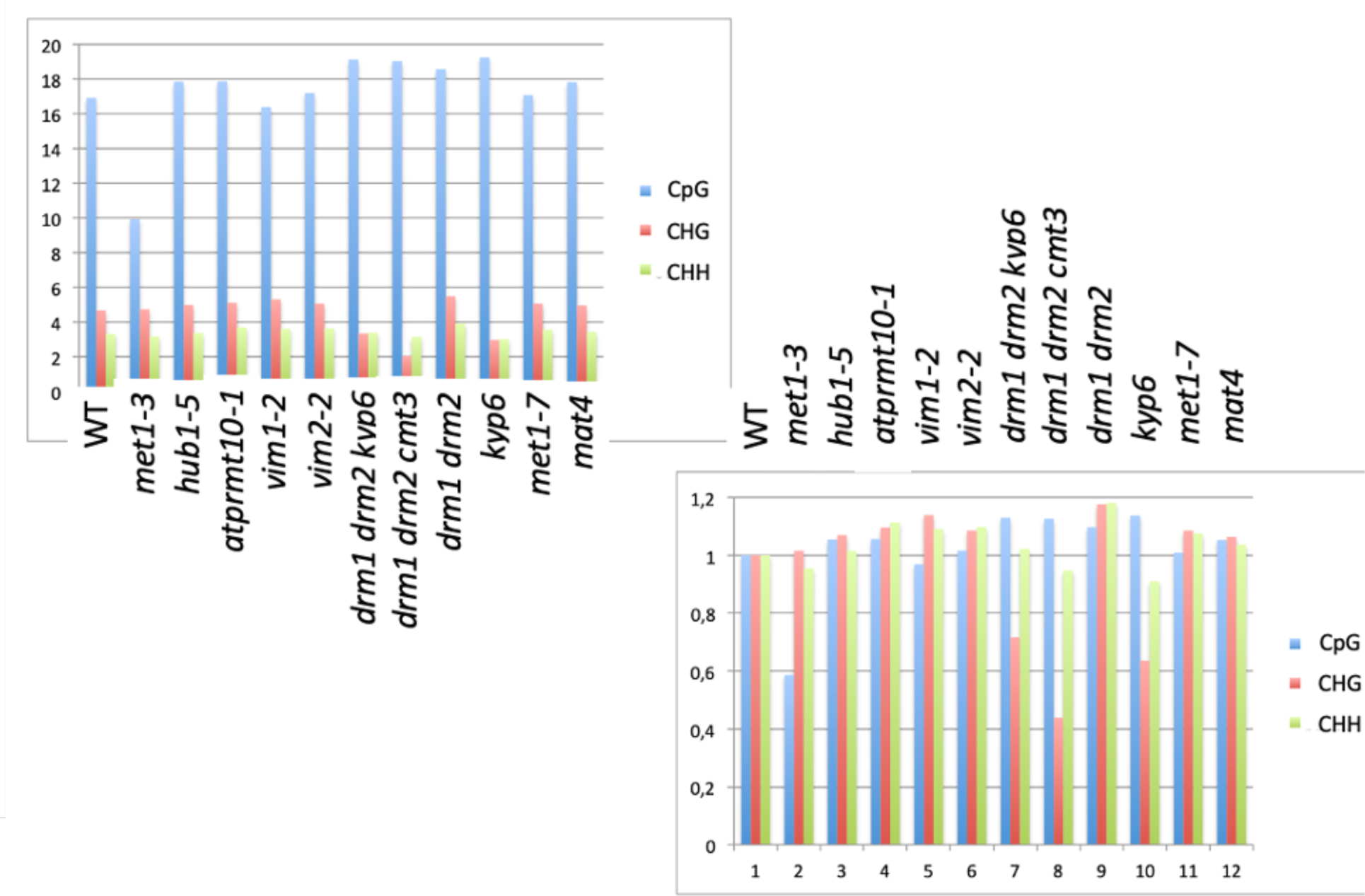
DNA methylation levels in CpG, CWG and CHH contexts for the WT and 11 mutants, in absolute numbers (Panel A) and normalised to the WT (Panel B).

**Figure 3:**
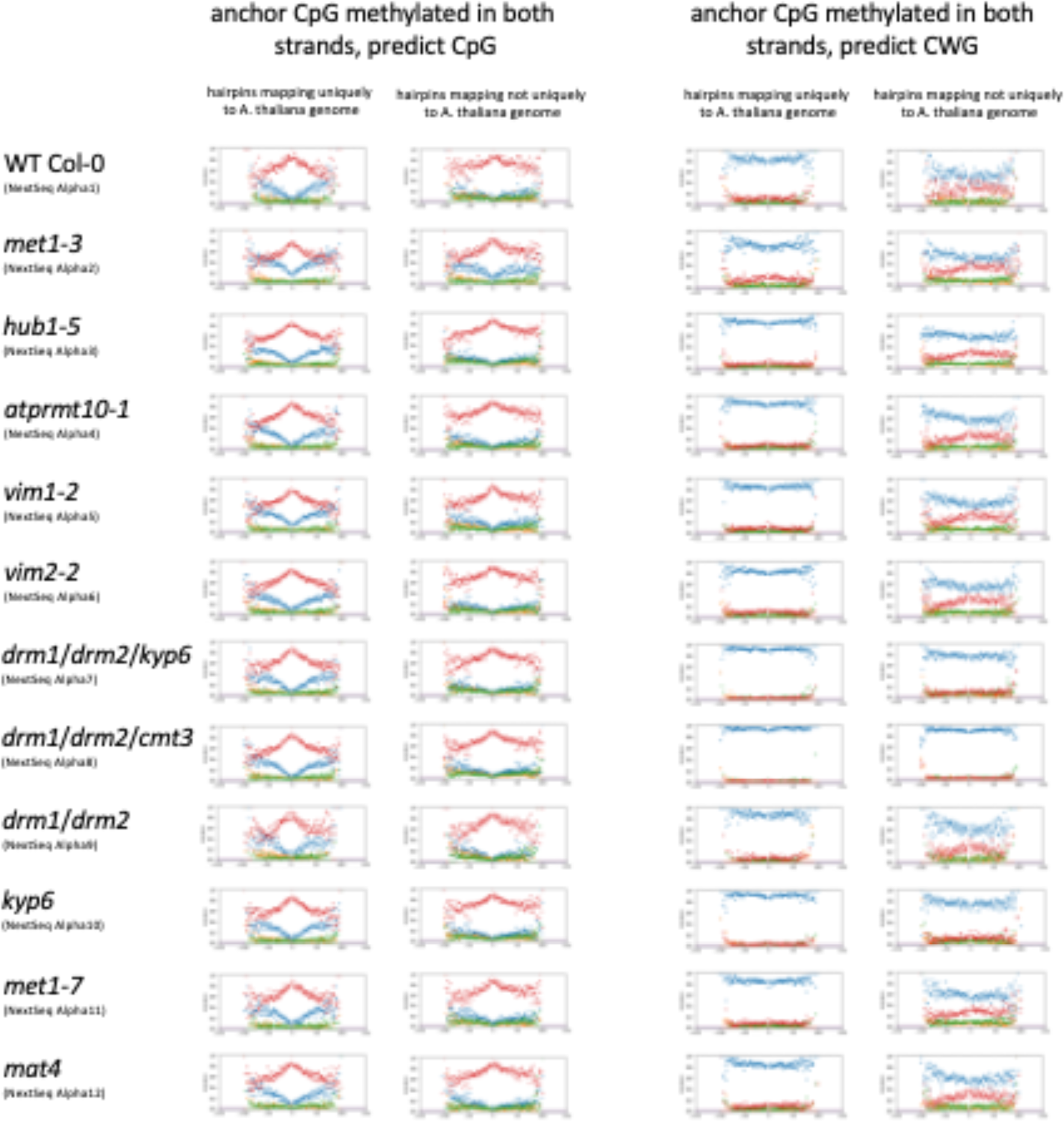
NextSeq DNA methylation predictability of CpG and CWG neighbouring to a methylated CpG site in WT plants and 11 mutants for reads that mapped uniquely or non-uniquely to *Arabidopsis* genomic DNA (WT panels repeated from Fig 1 are used for reference). Red dots represent fully methylated sites, blue indicate unmethylated sites and green and yellow signify hemimethylated sites.

CWG methylation is expected to be unaffected by impediments in MET1, but impacted when the CMT3/Kryptonite feed loop is disturbed. Compared to the WT reference (Fig. 4A), we observed a dramatic loss of CWG methylation predictability for mutants of CWG methylation enzymes (Fig. 4F), and also for *kyp6* and *cmt3* when combined with *drm1*/*drm2* (Fig. 4C/D) (drm1/drm2 alone was also analysed as a control Fig. 4E). The Met1 mutant (heterozygote *met1-3*) had no visible impact on the CWG predictability (Fig. 4B). The trends were similar for reads that mapped uniquely and those that did not map uniquely to the *Arabidopsis* genome (Fig. 4A-L)

**Figure 4:**
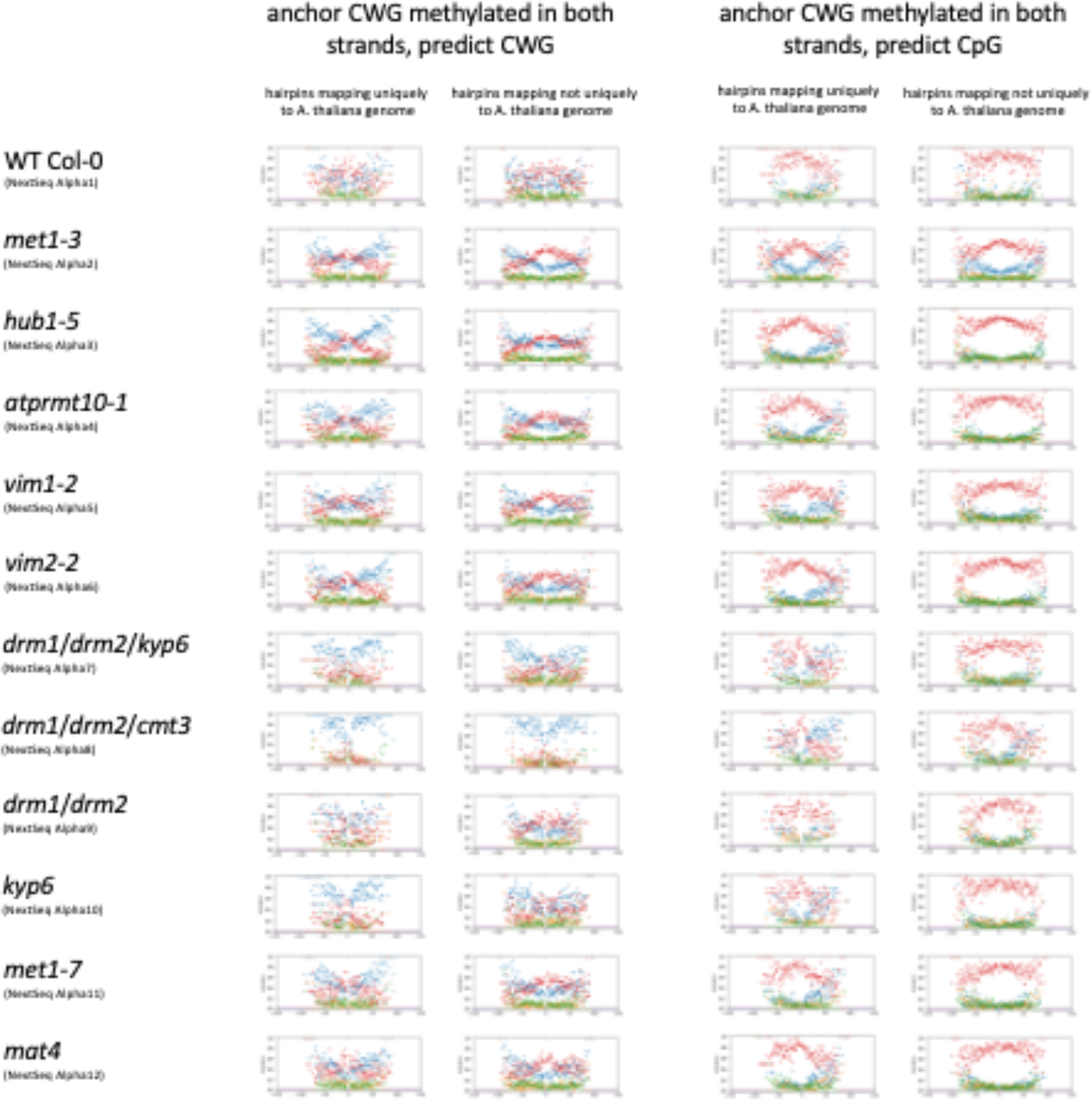
NextSeq DNA methylation predictability of CpG and CWG neighbouring to a methylated CWG site in WT plants and 11 mutants (WT panels are repeated from Fig. 1 and used for reference) for reads that mapped uniquely or not uniquely to *Arabidopsis* genomic DNA. Red dots represent fully methylated sites, blue indicate unmethylated sites, and green and yellow signify hemimethylated sites.

Across the mutants and experiments, the meta-analysis indicated that CpG methylation sites were are a good predictors of neighbouring CpG methylation but poor predictors of CWG methylation. In contrast, a fully-methylated CWG site was predictable for both neighbouring CWG methylation and CpG methylation.

### Fidelity factor calculation

Using in-house developed software and very high thresholds, the fidelity of the plant DNA methylation was then calculated. Disregarding non-conversion rates (calculated, for the CpG context, as 1% when analysing the chloroplast DNA, 0.3% for the spiked yeast DNA, and <0.27% for the hairpin cytosines), an ∼50× CpG fidelity factor was derived in the *met1-3* mutant, which was ∼30× for the WT plants. For CHG fidelity, the factor was approximately 15× for WT plants; therefore, the fidelity was considerably more accurate when all factors were present, and the methylated CpG sites in the *met1-3* mutant were replicated with higher fidelity than those of the WT.

### Conditional methylation probabilities for several *Arabidopsis* mutants estimated with MiSeq

To better understand the extension of the DNA conditional methylation predictability (lengths greater than one nucleosome) and to independently validate the original findings, an Illumina MiSeq sequencing was performed. MiSeq allows longer reads and, although it contains a price on throughput, it is generally more accurate (by virtue of a 4-channel system (4 dyes) compared to the 2-channel (2 dyes) system of NextSeq) than NextSeq. The original findings of conditional methylation probabilities were validated, although, despite the longer reads, the lengths were not sufficient to allow the DNA methylation predictability levels to decrease to the average levels. Therefore, the predictability extended to a distance greater than two nucleosomes (Figs. 5 and 6); however, the full scale of this extension is beyond the technical limits of the used Illumina platforms.

**Figure 5:**
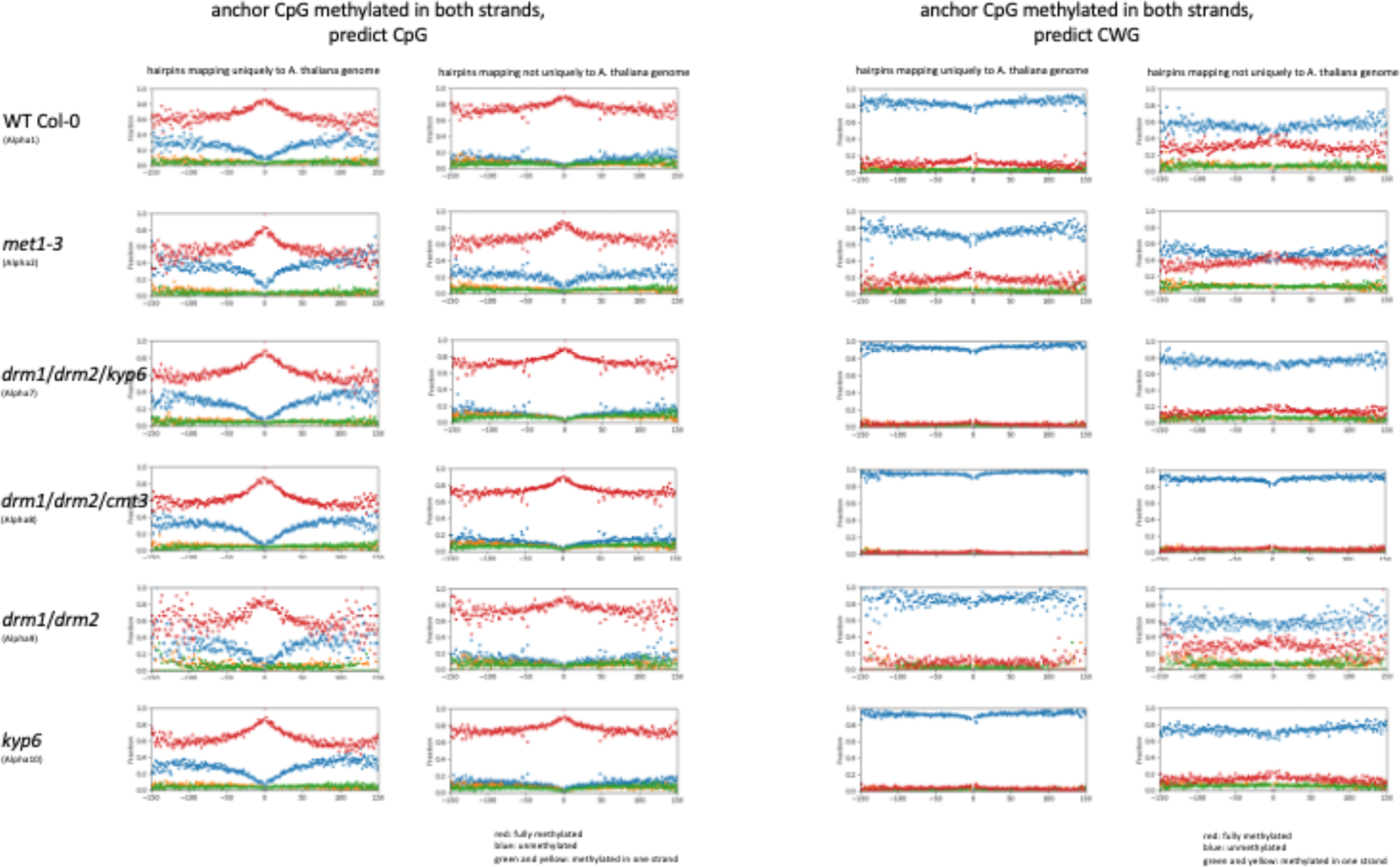
MiSeq DNA methylation predictability of CpG and CWG neighbouring to a methylated CpG site in WT plants and five mutants for reads that mapped uniquely and non-uniquely to *Arabidopsis* genomic DNA. Red dots represent fully methylated sites, blue indicate unmethylated sites, and green and yellow signify hemimethylated sites.

**Figure 6:**
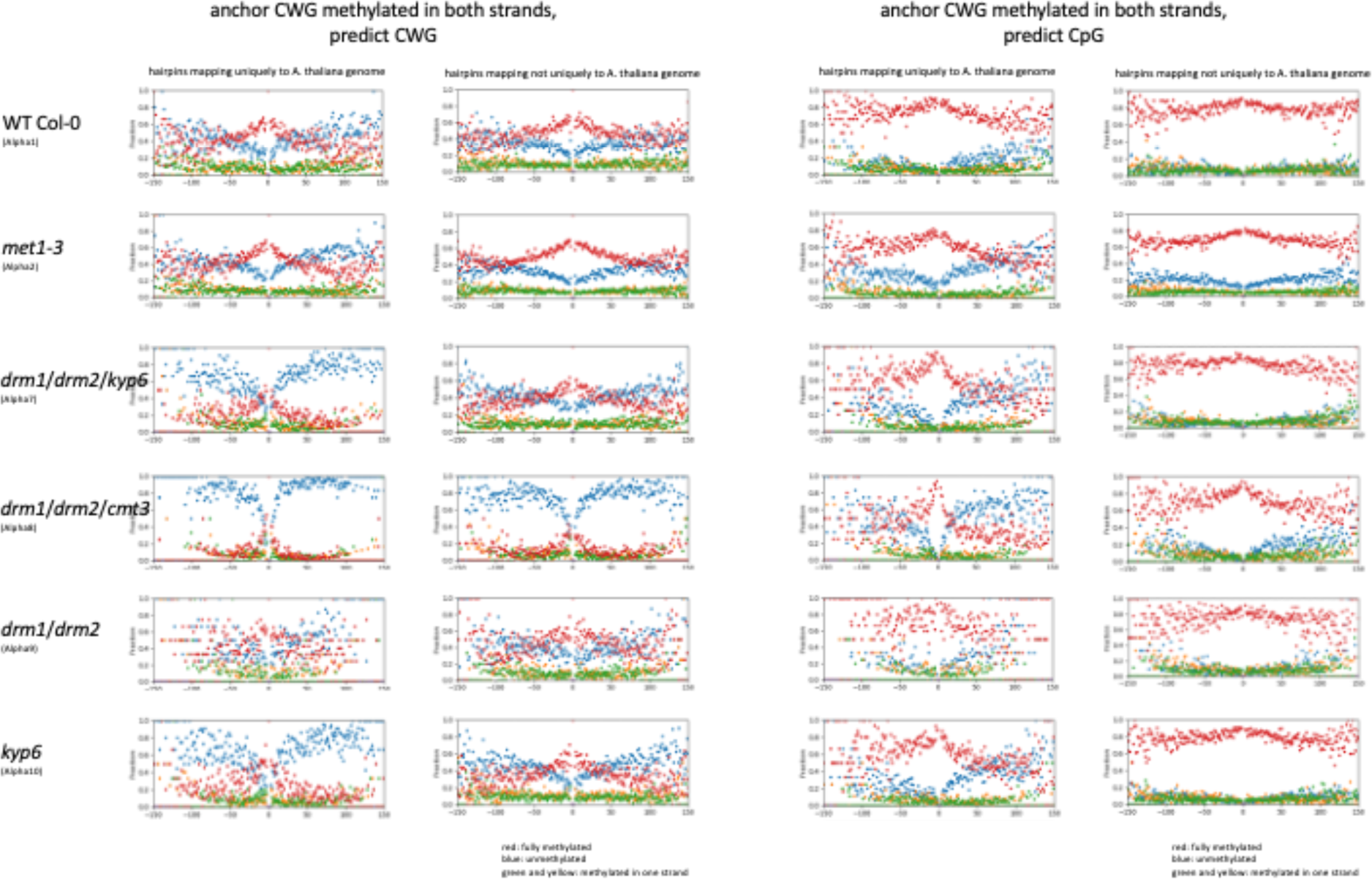
NextSeq DNA methylation predictability of CpG and CWG neighbouring to a methylated CWG site in WT plants and five mutants for reads that mapped uniquely and non-uniquely to *Arabidopsis* genomic DNA. Red dots represent fully methylated sites, blue indicate unmethylated sites, and green and yellow signify hemimethylated sites.

The meta-analysis validated the results obtained with NextSeq, with the CpG sites as good predictors of neighbouring CpG methylation (Fig. 5), while CWG methylation predicted both CpG and CWG neighbouring symmetric methylation (Fig. 6).

The improved accuracy (MiSeq vs NextSeq) allowed us to increase our analysis to include the predictability of neighbouring methylation of hemimethylated sites. CpG hemimethylation (Fig. 7) was not a good predictor of neighbouring full methylation (CpG or CWG), nor was CWG hemimethylation (Fig. 8), although both were good predictors for neighbouring hemimethylation. Note the increase at the central part of the plots for the hemimethylated sites (green and yellow), especially for CpG to CpG predictions (Fig. 7). Mirroring the activities at full methylation sites, including the anchoring and locking for neighbouring hemimethylation, CpG was a strong predictor of CpG, while CWG strongly predicted both CpG and CWG.

**Figure 7:**
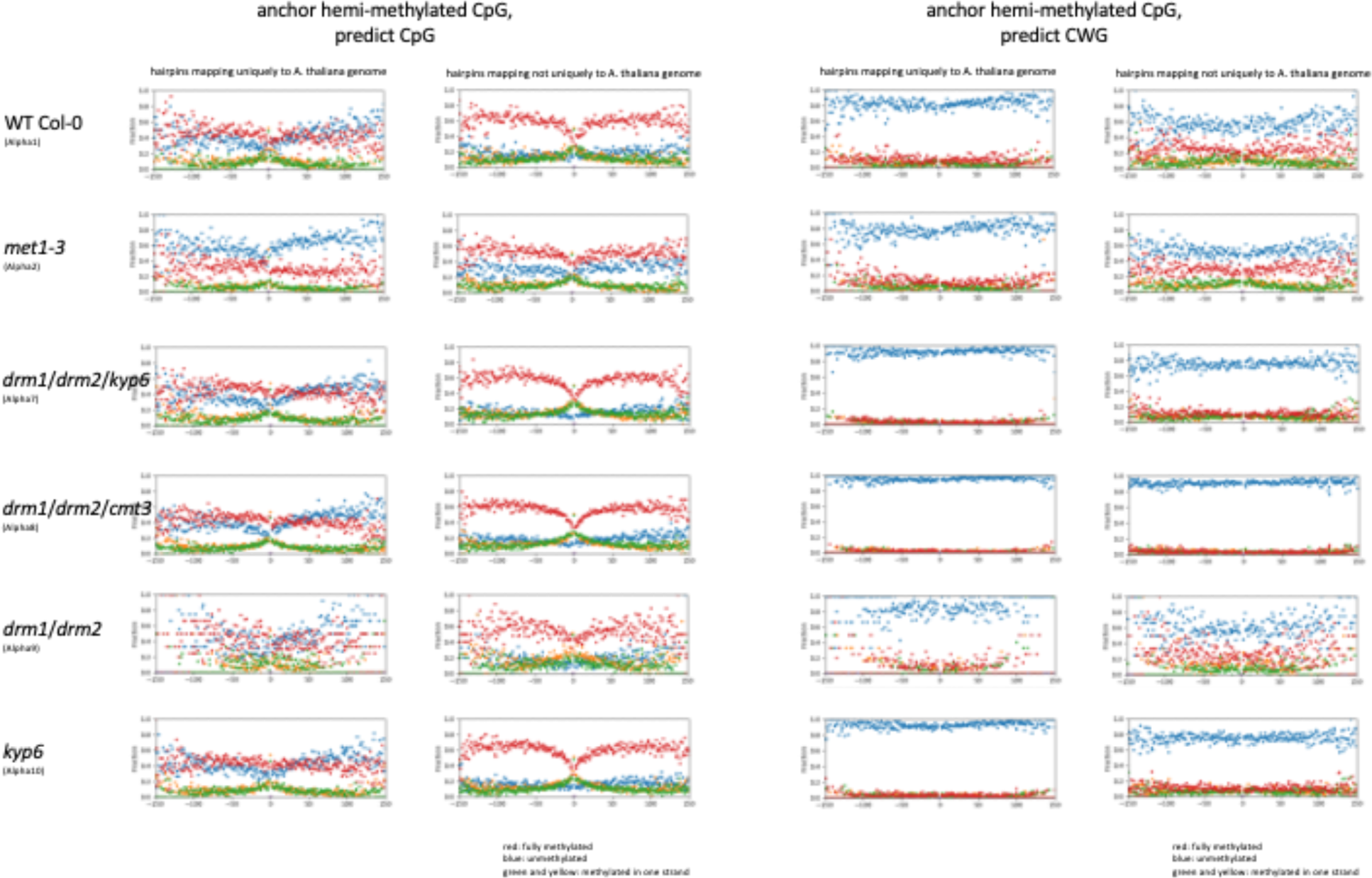
MiSeq DNA methylation predictability of CpG and CWG neighbouring to a hemimethylated CpG site in WT plants and five mutants for reads that mapped uniquely or not uniquely to *Arabidopsis* genomic DNA. Red dots represent fully methylated sites, blue indicate unmethylated sites, and green and yellow signify hemimethylated sites.

**Figure 8:**
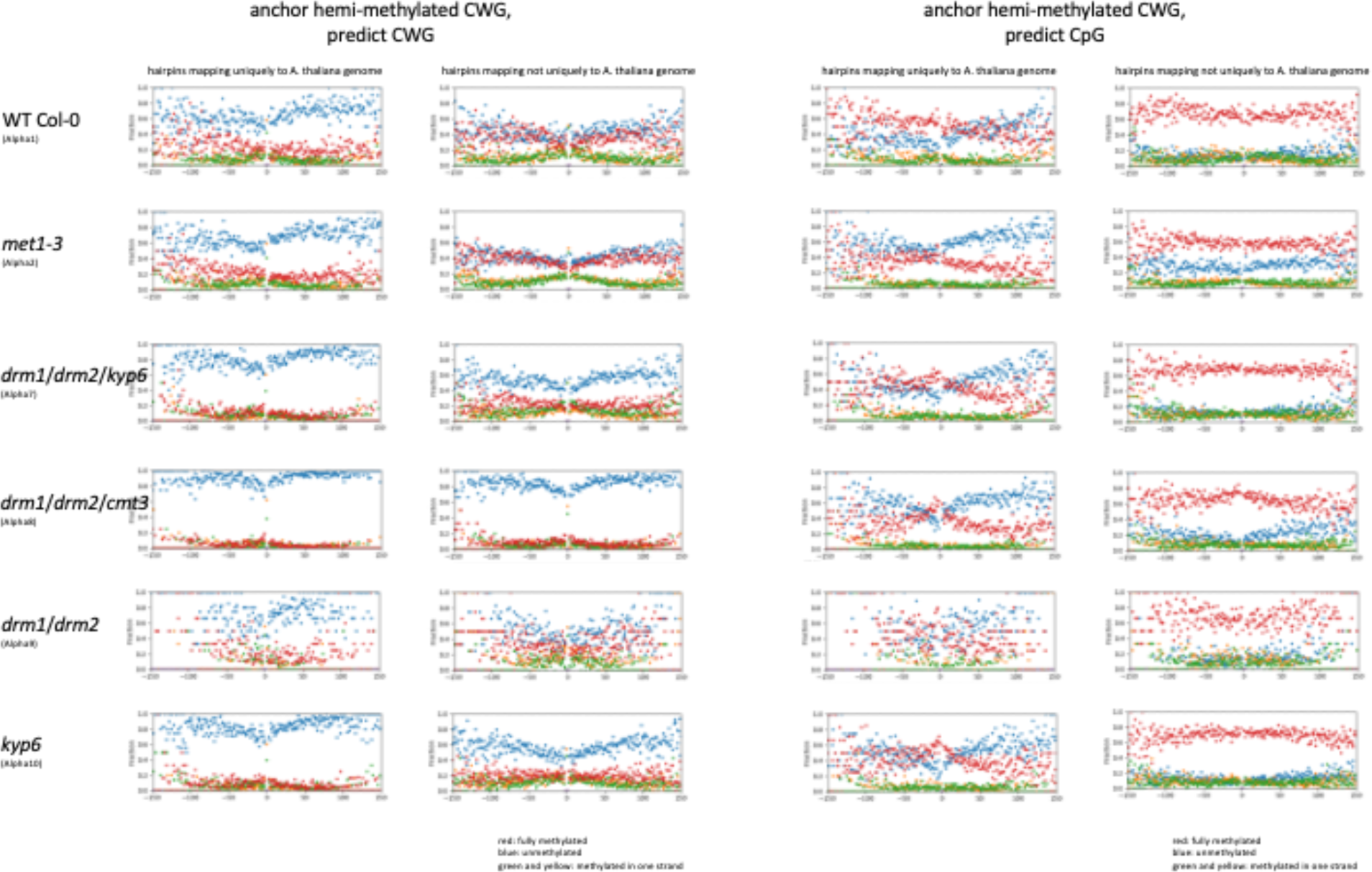
NextSeq DNA methylation predictability of CpG and CWG neighbouring to a hemimethylated CWG site in WT plants and five mutants, for reads that mapped uniquely or not uniquely to *Arabidopsis* genomic DNA. Red dots represent fully methylated sites, blue indicate unmethylated sites, and green and yellow signify hemimethylated sites.

### ‘Longitudinal’ long-range relationships in the same strand

The reversible terminator sequencing (Illumina) is technically limited to the range of a single DNA fragment molecule, and at extended distances, the anchor methylated CpG cue influence on neighbouring CpG methylation cannot be tested. However, the obtained results indicated that ‘longitudinal’ methylation predictability covers at least two nucleosome units. To collect the data for the average extent of the methylathion information influence, we used nanopore sequencing (Nanopore), which removes the hard limitation of read lengths. Methylation sensitive nanopore sequencing does not require bisulfite conversion which is intrinsically very damaging to long pieces of DNA (thanks to the perturbation of the ion flow by DNA modifications).

The *Arabidopsis* genome was fragmented into 8 kb pieces, and after testing our sample for 5mC levels (limited to CpG context), we determined a ∼15% false positive rate of the standard basecaller (the Guppy > MiniMap2 > Flappie pipeline) (Fig. S2). The calculation was based on the erroneous call of 5mC on unmethylated protoplast sequences in *Arabidopsis*; the same analysis revealed a 40% CpG methylation level for the nuclear genome (Fig. S2). The mitochondrial genome, despite the depletion of methylation, is not viable as a negative control because of the segmental duplication of a mitochondrial fragment in the centromere of chromosome 2 which is heavily methylated (thus the ∼35% CpG methylation present on our nanopore reads for mitochondrial DNA (Fig. S2). However, it offers another opportunity for internal controls, as reads that map to mitochondrial sequences should have two patterns of methylation: very low/no methylation if the DNA molecule originated from the organelle, or methylated if the DNA fragment originated from the nuclear genome (potentially, the methylation levels should be actually high, as the insertion on chromosome 2 is at the centromere region and thus heavily methylated). Mapping of the nanopore cumulative read distribution along the *Arabidopsis* genome revealed a sharp increase at the centromeres (Fig. S2), confirming its repetitive nature.

We applied the analysis of conditional methylation probabilities in *Arabidopsis* WT plants to the nanopore data and found that the CpG methylation probability on the same strand extended beyond the single nucleosome to a minimum of 4000 nucleotides for WT plants (the dataset had an ∼8.5× coverage) (Fig. 9). Interestingly, the correlation function exhibited characteristic ripples at a spacing of ∼180 base pairs (Fig. 9).

**Figure 9:**
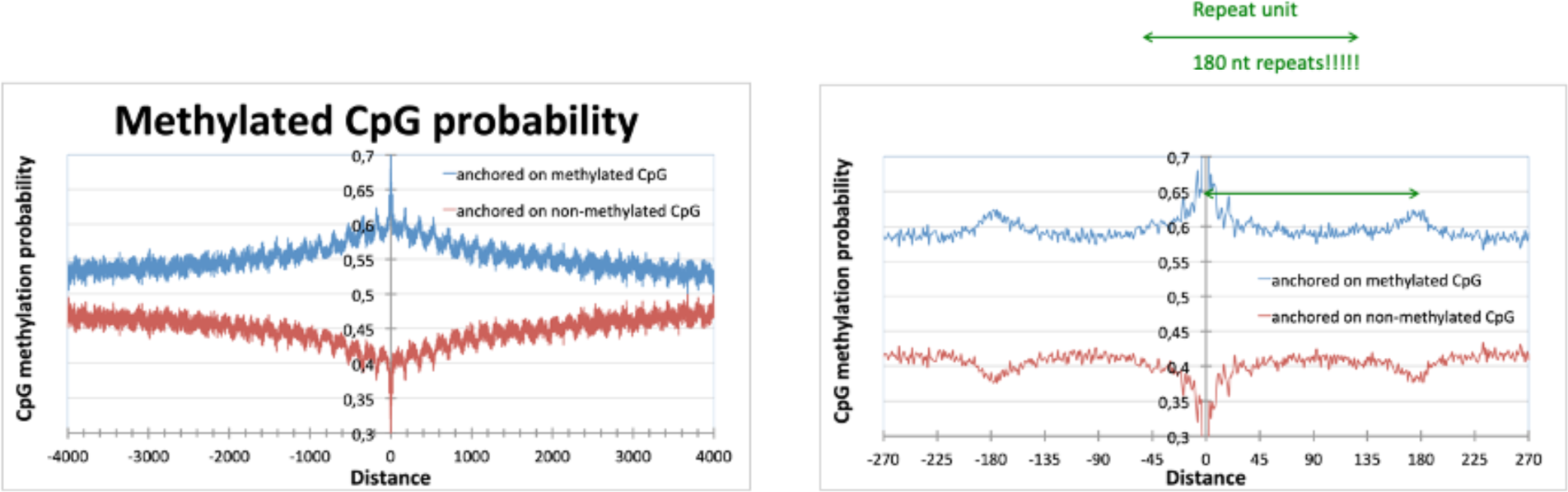
Long range correlation of CpG methylation (left panel), and zoom-in view highlighting the ripples at a spacing of ∼180 base pairs (right panel).

### ‘Longitudinal’ long-range relationships in the opposite strand

To test the relationships in the opposite strand, we used either hairpin bisulfite sequencing procedure or nanopore 1D^2^ chemistry which allows sequential sequencing of both strands of the same DNA fragment. Most consecutive sequences on individual pores were from different fragments, but ∼10% were hairpin-mimicking results that corresponded to approximately 7× coverage of the *Arabidopsis* genome. The remaining data (approx. 60× coverage) were valid and useful standard (1D) reads.

## Discussion

A study of DNA fidelity in DNA replication needs to address the degree of methylation status maintenance of the two daughter strands upon cell division. The superior method for identifying methylated cytosines in DNA is the use of bisulfite sequencing. To precisely read the methylation levels onto pairing strands, the strands need to be linked with a hairpin loop (single-cell epigenetic sequencing is not developed enough to allow for a fidelity study). This strategy is ideal for measuring fidelity in both a CpG and CWG context; however, it is unsuitable for fidelity testing at CHH sites. Transgenerational studies are required in these situations, since CHH has a non-symmetrical methylation in which the methylation information is recorded on only one of the strands, and the newly synthesised strands must receive cues from areas other than the counter-strand. For this study, we used a biotinylated hairpin to link paired strands prior to BS treatment, with a strategy to enrich sequences with paired strands using by affinity pull-downs.

In addition to the writer domain (the part of the protein that executes the DNA methylation step), DNA MTases have numerous reader domains that help guide the protein to the appropriate locations within the chromatin, while enzymes that deposit marks in the chromatin read the DNA methylation status. Many examples of these processes occur in eukaryote epigenetics, and the literature provides extensive information regarding this subject. A clear example is that between the pair CMT3 and kryptonite in plants and their self-reinforcing loop. In this case, CMT3 has a DNA methyltransferase domain as well as BAH and a Chromo domain which are responsible for binding methylated Histone H3 Lysine 9 (H3K9me2) and guiding CWG methylation to the sites of H3K9me2. Kryptonite, in turn, in addition to its SET domain responsible for direct methylation of H3K9, harbours an SRA domain that is important for recognising methylated DNA and guiding kryptonite-driven histone methylation to sites of CWG methylation. For this study, we selected a range of mutants that had impaired DNA methylation, histone methylation, and ubiquitination modification capacity or ability to read these marks. We used mutants that had the same genetic background ecotype (Columbia) and isolated DNA from the same tissues and age groups. As expected, we observed a decrease of CpG methylation with the *met1-3* mutant and a decrease of CWG with *kyp6* or *cmt3* mutant backgrounds.

Extensive research has been conducted to identify links among the multiple pathways that regulate DNA methylation and the levels and genomic positions of cytosine methylation in different contexts. Here, we extended the analysis to investigate the links between DNA methylation and the probability of the methylation of neighbouring cytosines being methylated in the vicinity of previously methylated cytosines. We adopted the assumption that reads that map only once to one of the five *Arabidopsis* chromosomes were euchromatin regions, and those that could not uniquely be mapped were locked in the heterochromatin. This postulation is supported by the CpG methylation levels, which were higher for the heterochromatin in our bisulfite data, and by the copy number variation, which showed a sharp increase at the centromeres of the cumulative read distribution along the full size of the five *Arabidopsis thaliana* chromosomes (but not at plastids or mitochondria). The sharp increase at the centromeres also suggests that the *A. thaliana* genome annotation we used may be missing repetitive regions near the centromeres.

Our analysis demonstrated a clear link between a symmetric methylated site and the probability that a nearby, similar site, will also be methylated. Moreover, the data also show a level of hierarchy, with the CpG probability being higher than the global average in the vicinity of methylation at CpG and at CWG sites, while the CWG methylation was higher only in the vicinity of the CWG but not the CpG sites. Our initial meta-analysis indicated a slight decrease of methylation probability near sites of methylated DNA, but the shot reads generated with NextSeq did not allow us to determine the extent of this probability along the DNA. The initial test with MiSeq indicated a minimum length of one nucleosome, although, considering the distance, the probability was much higher than the global average. We then applied nanopore technology, which generates very long reads, and determined that the such probability remained higher than the global average at a distance exceeding 4000 nt. With the decay of probability, ripples were observed at a spacing of ∼180 base pairs, which could be due to nucleosome and/or centromere repeats. We then ruled out or confirmed the latter by reprocessing the data-removing sequence that mapped to the centromere regions.

## Conclusion

The fidelity of DNA methylation is crucial for the proper functioning of cells since DNA methylation is necessary for cell differentiation and organism development. The fidelity of this process is crucial for proper gene regulation, cellular function, and other biological processes, in addition to playing a fundamental role in inheritance. MTases are aided by a series of cues on the chromatin, whereby the integrity of its enzymatic activity increases with the presence of supporting epigenetic machinery. Mutant plants with impaired MET1 or CMT3 activities had lower levels of methylation at the CpG and CWG sites, respectively, although the remaining sites were maintained with higher degrees of fidelity than those of WT plants. In addition, the results of this study indicated that CpG and CWG methylated sites were good predictors of similar methylation at neighbouring sites and that these predictor factors extended to a length greater than that of a single nucleosome indicating a long range influence.

## Materials and methods

### Plant materials (for hairpin)

WT Col-0 and mutants seeds were surface chlorine sterilised (Vapor-Phase sterilisation), stratified for four days at 4°C, and grown in soil in a greenhouse for 22 days. Arial parts of the seedlings were collected, flash-frozen in liquid nitrogen and stored at −80°C until processing. The mutants used in this study were T-DNA insertions and were acquired from TAIR seed stock. The mutants use are *met1-3* (CS16394), *met1-7* (SALK_076522), *vim1-2* (SALK_050903), *vim2-2* (SALK_133677), *drm1*/*drm2*/*kyp6* (CS16388), *kyp-6* (CS16385), *drm1*/*drm2*/*cmt3* (CS16384), *hub1-5* (SALK_044415C), *atprmt10-1* (SALK_047046C), *mat4* (SALK_011935) and *drm1*/*drm2* (CS16383).

The presence of T-DNA insertions was confirmed by genotyping the different plant lines using a T-DNA left border and allele-specific oligonucleotides (Oligonucleotide sequences are listed in Table 1)

**Primer table 1.**
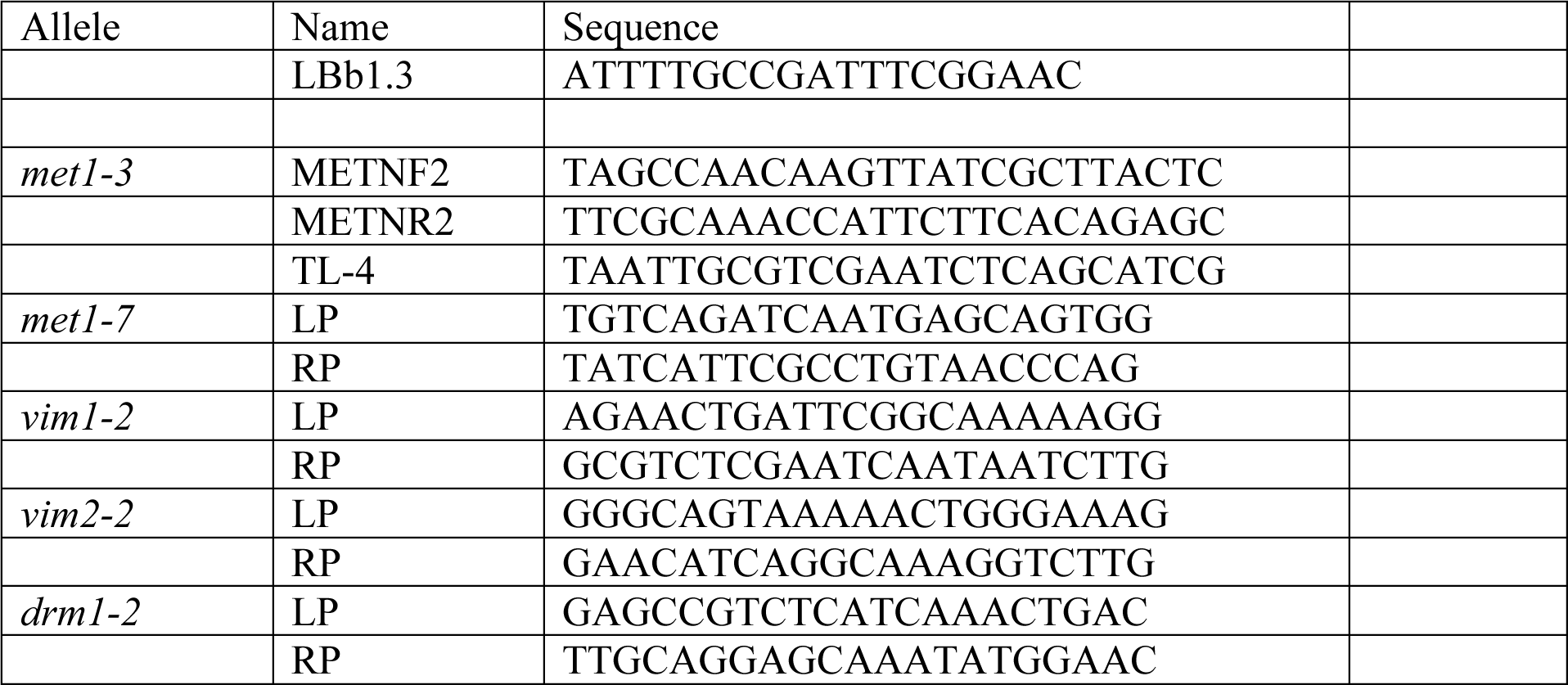
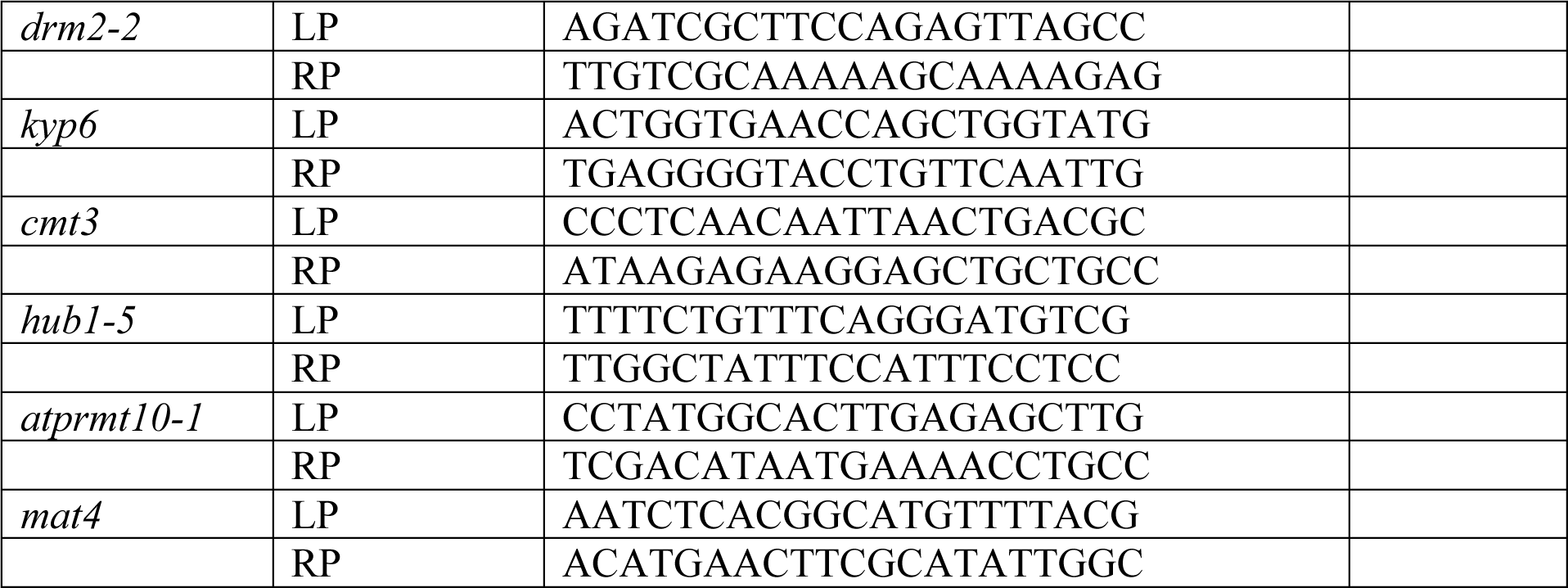

### Hairpin library preparation

*Arabidopsis thaliana* genomic DNA was extracted using the DNeasy Plant Mini Kit (Qiagen). One microgram of *Arabidopsis* genomic DNA was spiked with 1% unmethylated *S. cerevisiae* genomic DNA (Merck) and sonicated to 200-bp fragments using a Covaris M220. The fragments were subjected to end repair and dA tailing using the KAPA End Repair & A-Tailing Enzyme (KapaBiosystems). A biotin-modified hairpin (P-CGCCGGCGGCAAG/iBiodT/GAAGCCGCCGGCGT) (Sigma) was annealed by fast cooling/dilution in ice-cold water for 10 minutes at 90°C, and Illumina TruSeq adapters (Illumina) were ligated to the fragments (molar ratio 29:1:1) overnight using the KAPA DNA Ligase Enzyme (KapaBiosystems). After purification with AMPure XP beads (Agencourt; bead to probe ratio 0.8:1), DNA fragments were pulled down using Dynabeads MyOne Streptavidin C1 beads (Invitrogen). To release the DNA from the beads, 20 μg of Proteinase K (GeneON) was added, and the slur was incubated overnight at 50°C, followed by 10 minutes of inactivation at 80°C. The cleared DNA was subjected to bisulfite conversion using the EZ DNA Methylation-Gold kit (ZymoResearch) and purified with AMPure XP beads (Agencourt; beads to probe ratio 5:1). The libraries were prepared using ∼ 30 ng of the single-stranded uracil-containing DNA as templates, amplified with KAPA HiFi HS Uracil+ RM (KapaBiosystems) using KAPA Library Amplification Primer Mix (KapaBiosystems), and purified with AMPure XP beads (Agencourt; beads to probe ratio 0.9:1). Quality control and concentration determination were performed using High Sensitivity D1000 ScreenTape (Agilent) and qPCR. The libraries were normalised, pooled, and diluted for sequencing. Sequencing was performed using NextSeq 500 and MiSeq sequencers (Illumina) following the manufacturers’ protocols.

### Data analysis

A NextSeq data set (with the different backgrounds anonymised) was first screened for the presence of the hairpin, the condition necessary to test DNA methylation fidelity, and then for non-conversion rates. This was conducted robustly by assessing yeast gDNA that was spiked-in, chloroplast sequences (non-methylated in plants) and the hairpin sequence (synthesised with standard cytosines (i.e. non-methylated cytosines). The sequence results were subsequently analysed for the level and context of methylation. The results indicated that different backgrounds displayed different levels in particular contexts, as expected. At this point (and only at this point), the anonymisation of the backgrounds was removed, and the global levels for the various contexts fit the expectations for the respective backgrounds sequenced. For example, a decrease of methylation was observed in the CpG context when MET1 mutants were used or a decrease of methylation occurred in the CHG context when CMT3 or kryptonite mutants were sequenced. The MiSeq was analysed in a similar fashion, although the samples were not anonymised.

### ONT library preparation

*Arabidopsis thaliana* genomic DNA was extracted using the DNeasy Plant Mini Kit (Qiagen). WT plants were used, along with the mutants: *met1-7*, *kyp6*, *drm1/drm2/kpy6*, *drm1/drm2/cmt3* and *drm1/drm2*. Between 0.9 and 1.8 μg of each type of genomic DNA was processed by DNA shearing by applying 32 μL to a g-TUBE (Covaries) and centrifuging at 13523 *g* for 2 × 1 minute to obtain ∼ 8 kb fragment sizes. The BluePippin system (Sage science) with 0.75% dye-free Agarose cassettes and S1 markers was used to select fragments >6 kb.

For the nanopore 1D sequencing, the purified and size selected DNA was prepared for barcoding (EXP-NBD104, ONT, Oxford, UK) and sequencing with the Ligation Sequencing kit 1D (SQK-LSK109, ONT, Oxford, UK) sequencing kit protocol. Briefly, between 0.9 and 1.8 μg of each type of genomic DNA was repaired and end-prepped with NEBNext FFPE DNA Repair Mix and Ultra II End-prep enzyme mix (NEB) for 5 minutes at 20°C followed by 5 minutes at 65°C for inactivation of the enzymes, and purified with 1X AMPure XP beads (Beckman Coulter). The genomic DNA was barcoded, adding Native Barcoding Expansion (ONT, Oxford, UK) and Blunt/TA ligase mix (NEB) was added for 10 minutes at room temperature and purified with 1X AMPure XP beads (Beckman Coulter). Adapter Mix II (ONT, Oxford, UK) and Quick T4 DNA Ligase (NEB) were added and incubated for 10 minutes at room temperature and purified with 0.5X AMPure XP beads (Beckman Coulter), and Long Fragment buffer (ONT, Oxford, UK) was used to purify the library molecules, which were recovered in an Elution buffer (ONT, Oxford, UK) [concentration of the polled DNA: 20.2 ng/μL]; quality control was conducted with High Sensitivity D1000 ScreenTape (Agilent)] by incubating the beads for 10 minutes at 37°C. The purified barcoded libraries were combined with the Sequencing Buffer and Loading Beads (ONT, Oxford, UK) and loaded onto a primed R9.4 Spot-On Flow cell. The flow cell was mounted on a MinION Mk 1B device (ONT) for sequencing with MinKNOW versions 1.1.15-1.1.21 with the NC_48Hr_Sequencing_Run_FLO-MIN106-LSK109 script.

For nanopore 1D^2^ sequencing, *Arabidopsis thaliana* WT Col-0 genomic DNA was extracted using the DNeasy Plant Mini Kit (Qiagen). Approximately 2 μg of sheared genomic DNA was repaired and end-prepped with NEBNext FFPE DNA Repair Mix and Ultra II End-prep enzyme mix (NEB) for 15 minutes at 20°C followed by 5 minutes at 65°C for inactivation of the enzymes and purified with 1X AMPure XP beads (Beckman Coulter); 1D^2^ Adapters (ONT, Oxford, UK) and Quick T4 DNA Ligase (NEB) were added and incubated for 10 minutes at room temperature and purified with 0.4X AMPure XP beads (Beckman Coulter). 1D^2^ Adapters (ONT, Oxford, UK) and Quick T4 DNA Ligase (NEB) were added and incubated for 10 minutes at room temperature and purified with 0.4X AMPure XP beads (Beckman Coulter). Adapter Mix II (ONT, Oxford, UK), ONT Ligation Buffer (ONT, Oxford, UK) and Quick T4 DNA Ligase (NEB) were added and incubated for 10 minutes at room temperature and purified with 0.4X AMPure XP beads (Beckman Coulter) using Long Fragment buffer (ONT, Oxford, UK) been used to purify the library molecules, which were recovered in Elution buffer (ONT, Oxford, UK) by incubating the beads for 10 minutes at 37°C. The purified library was combined with the Sequencing Buffer and Loading Beads (ONT, Oxford, UK) and loaded onto a primed R9.5 Spot-On Flow cell. The flow cell was mounted on a MinION Mk 1B device (ONT) for sequencing with MinKNOW versions 1.1.15-1.1.21 with the NC_48Hr_Sequencing_Run_FLO-MIN106-LSK109 script.

## Acknowledgements

We are grateful to the green house facility of the Institute of Biochemistry and Biophysics for support on Arabidopsis handling.

