## Supplementary material for "Longitudinal conditional probability of symmetric DNA methylation in *Arabidopsis* plants": Fig. S

FigS1a1

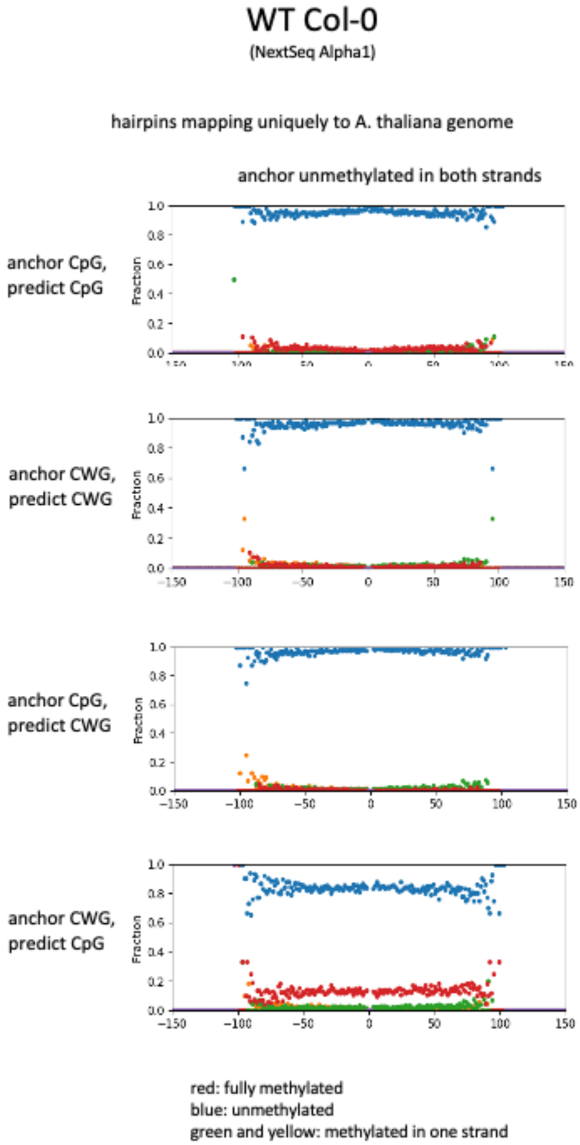


FigS1a2

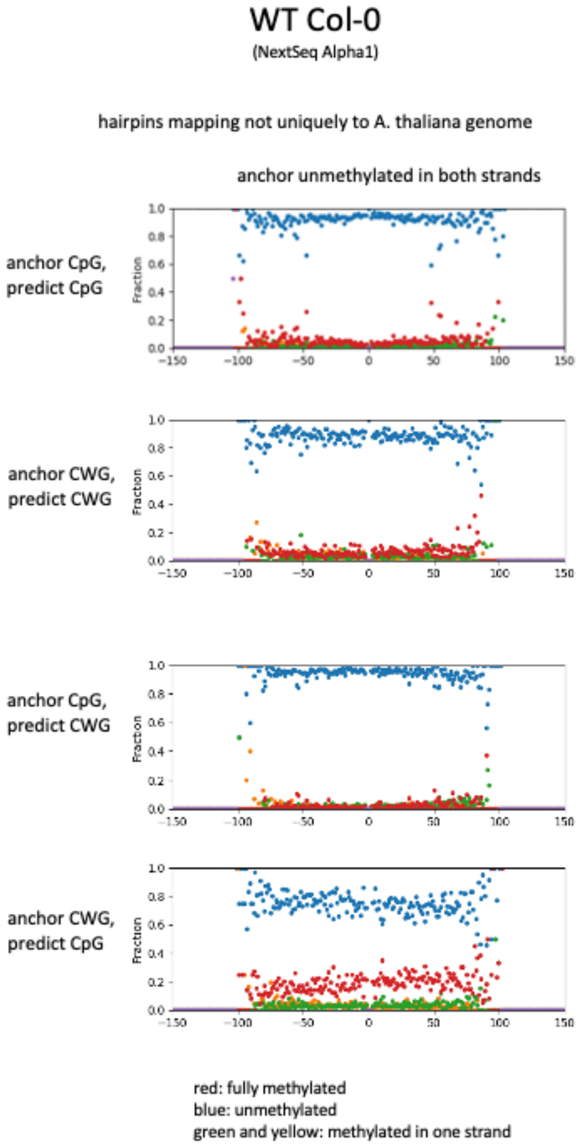


Figure S1: DNA methylation predictability of CpG and CWG neighbouring to an unmethylated CpG or CWG site in WT plants for reads that mapped uniquely (S1a1) or not uniquely (S1a2) to *Arabidopsis* genomic DNA. Red dots represent fully methylated sites, blue indicate unmethylated sites and green and yellow signify hemimethylated sites.

A)

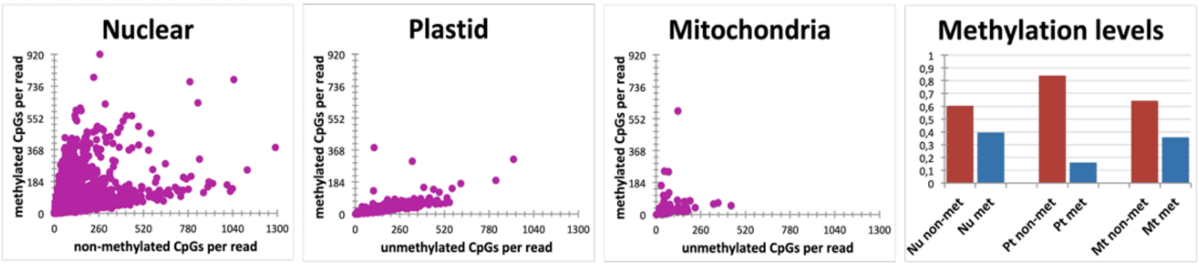


B)

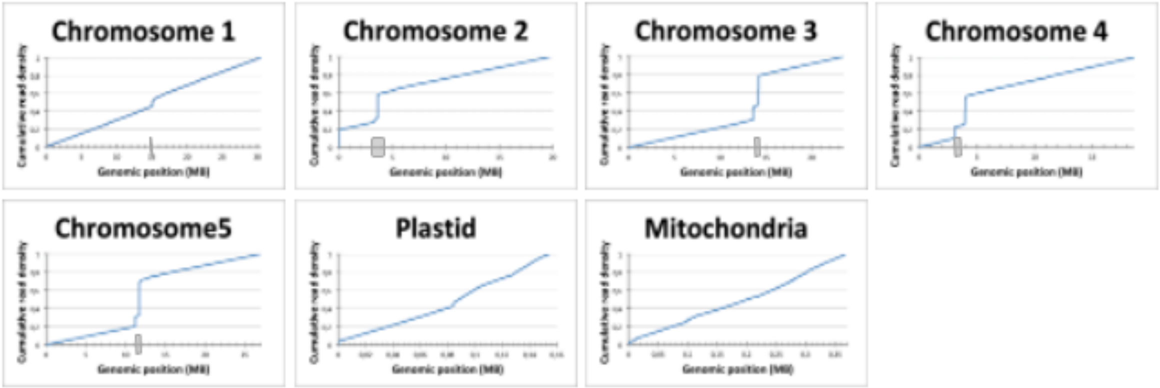


Figure S2: Nanopore CpG methylation validation. (A) 5mC base calling by Flappie was mapped by Minimap2 and grouped according to the source of origin (Nuclear, Plastid, Mitochondria), with the total percentage of methylated and unmethylated cytosines displayed as a bar plot. (B) Cumulative read distribution along the full size of the five Arabidopsis thaliana chromosomes. The sharp increase at the centromeres suggests that the current A. thaliana genome annotation may be missing repetitive regions near centromeres. Centromeres are marked with grey boxes in their genomic positions.
